# Emergence of the Novel Infectious bursal disease viruse Variant in Vaccinated Poultry Flocks in Egypt

**DOI:** 10.1101/2023.11.13.566865

**Authors:** Momtaz A. Shahein, Hesham A. Sultan, Ali Zanaty, Amany Adel, Zienab Mosaad, Dalia Said, Ahmed Erfan, Mohamed Samy, Abdullah Selim, Karim Selim, Mahmoud M. Naguib, Heba Hassan, Osama El Shazly, Zeinab A. El-badiea, Mahmoud K. Moawad, Abdelhafez Samir, Mohamed El Shahaby, Eman Farghaly, Samah Eid, Mohamed N Abdelaziz, Mohamed M Hamoud, Osama Mehana, Naglaa M. Hagag, Ahmed Samy

## Abstract

Infectious bursal disease viruses (IBDVs) have a profound impact on poultry production worldwide, directly causing mortality rates of up to 100%, and indirectly through their immunosuppressive effects. Since the emergence of the antigenically modified very virulent IBDV (vvIBDV) in Egypt in late 1999, the country has experienced recurrent outbreaks with high mortality rates and typical vvIBDV gross lesions. However, a notable shift occurred in 2023, characterized by a substantial increase in reported subclinical IBDV cases exhibiting atrophied bursa and associated immunosuppression. To assess the field situation, we examined samples from 21 farms in 2023 and 18 farms from 2021 and 2022, all of which experienced IBD outbreaks based on clinical diagnosis. These samples were submitted to our laboratory for confirmatory testing and subsequently subjected to VP2-HVR sequencing. Phylogenetic analysis revealed that all samples collected in 2021 and 2022 clustered with classical virulent strains and very virulent IBDV. In 2023, one sample clustered with the Egyptian vvIBDV, while one sample clustered with classic virulent IBDV, and the remaining 2023 samples clustered with the Chinese novel variant IBDV (nVarIBDV). The alignment of deduced amino acid sequences for VP2 revealed that all Egyptian classic virulent strains were similar to the Winterfield or Leukert strains. In contrast, vvIBDV strains exhibited two out of the three typical residues found in Egyptian antigenically atypical vvIBDV, namely Y220F and G254S, but not A321T, and one sample was identical to the European vvIBDV (emerged in 1989). Meanwhile, all variant strains recognized in the present study exhibited typical residues found in variant IBDV, in addition to the three conserved amino acid residues found only in Chinese variant IBDVs. However, all Egyptian variant strains showed a mutation at position 321 (321V), which represents the most exposed part of the capsid and is known to have a massive impact on IBDV antigenicity, with the exception of one sample that had 318G instead. This report highlights the emergence of a new variant IBDV clustered with the Chinese new variant in Egypt, causing bursa atrophy and spreading subclinically in broiler farms over a wide geographic distance, resulting in massive economic losses due to immunosuppression.

## Introduction

Infectious bursal disease viruses (IBDVs) pose a severe threat to global poultry production, with certain strains causing mortality rates as high as 100% (Eterradossi et al. 1992; Eterradossi and Saif 2020). IBDVs primarily target IgM-bearing B-lymphocytes within the bursa of Fabricius (Burkhardt and Müller 1987; Müller 1986) Subsequently, the virus spreads to other lymphatic organs, leading to either permanent or transient immunosuppression. This immunosuppression increases susceptibility to other pathogens and diminishes the efficacy of commonly used vaccines.(Eterradossi and Saif 2020).

Infectious bursal disease (IBD) is caused by the infectious bursal disease virus (IBDV), a highly contagious, non-enveloped, bi-segmented RNA virus that belongs to the Avibirnavirus genus of the Birnaviridae family (Müller, Islam, and Raue 2003). The IBDV genome consists of two segments, A and B. Segment A encodes four viral proteins, including two structural proteins, VP2 and VP3, the viral protease VP4, and the nonstructural protein VP5, while segment B only encodes VP1, the RNA-dependent RNA polymerase (Müller and Becht 1982; Raja et al. 2016). VP2 forms the outer capsid protein and is associated with virulence, cell tropism, and represents the major target of protective humoral immunity (Jackwood et al. 2008). VP2 is composed of the base (B), shell (S), and projection (P) domains. The P domain forms the outer part of the protein and contains four loops named P_BC_, P_HI_, P_FG_, and P_DE_. The major immunogenic epitopes within the P domain are located in the hypervariable region (HVR), which spans from amino acid positions 206 to 350. Within the HVR, there are two major hydrophilic peaks, A (212–224 aa) and B (312–324 aa), separated by two hydrophobic peaks 1 and 2, 248–254 and 279–290. These peaks, particularly major peaks A and B, are responsible for variations in IBDV antigenicity (Vakharia et al. 1994).

Only isolates belongs to serotype 1 are pathogenic to chickens that classified based on antigenicity and pathogenicity in to classical strains (cIBDV), antigenic variant strains (varIBDV), very virulent strains (vvIBDV), (Van den Berg et al. 2004; Eterradossi and Saif 2020), also can include attenuated IBDV, and novel variant IBDV (nVarIBDV) (Gao et al. 2023). Classical IBDV (cIBDV), which first detected in 1957 in Gumboro, Delaware, USA, causing mortality rates ranging from 1% to 50% (Cosgrove 1962) and immunosuppression that impacts the host’s immune response and the effectiveness of other vaccinations (Müller, Islam, and Raue 2003). Variant IBDV (varIBDV) was first reported in the late 1980s that is different antigenically from cIBDV and can evade neutralising antibodies produced by vaccination against the cIBDV. It is characterized by little to no mortality with no or little clinical signs but significant bursal and splenic damage (Eterradossi et al. 1997). A novel Variant IBDV (nVarIBDV) emerged in China since 2015, displaying significant genetic distinctions from earlier varIBDV strains but still associated with subclinical infections in chickens, making it potentially easy to overlook. Nevertheless, it can lead to marked bursal atrophy and subsequent immunosuppression that associated with massive economic losses (Huang et al. 2021; Wang et al. 2022). Very virulent IBDV (vvIBDV) emerged in Europe, representing the acute form of IBDV. It is characterized by a high mortality rate of 50-100%, high spread rate, typical clinical signs, even in the presence of maternal antibodies. vvIBDV affects immune organs beyond the bursa, resulting in sever reduction of immune response (Huang et al. 2021).

The IBDV recorded for the first time in Egypt in Egypt by El-Sergany and colleagues in 1974 (Hassan 2004). By the late 1980s, the European form of very virulent IBDV (vvIBDV) had spread to Egypt, as reported by El-Batrawi in 1990 (Zierenberg et al. 2000; El-Batrawy 1990). This strain caused acute outbreaks, even in vaccinated chickens, including native and foreign breeds, immunized with both live and inactivated classic vaccines (Hassan 2004; Hassan, Afify, and Aly 2002). In 1999, an antigenically atypical vvIBDV strain emerged, characterized by specific mutations (Y220F, G254S, and A321T) (Eterradossi et al. 2004). These mutations had a significant impact on the virus’s antigenicity, leading to a loss of binding with 6 out of 8 neutralizing anti-VP2 monoclonal antibodies that were originally raised against the classical strain. In contrast, the European vvIBDV that loss binding to 2 out of 8 antibodies (Eterradossi et al. 2004; Samy et al. 2020). Despite intensive vaccination efforts, mainly based on a live-attenuated, intermediate plus, classical strain-based vaccine, that shown to confer experimental clinical protection (Eterradossi et al. 2004), the antigenically variant vvIBDV strain continued to circulate in Egypt, with some isolates retaining the Ala_!Il_Thr mutation at position 321, enabling them to maintain binding activity with 5 out of 8 neutralizing anti-VP2 monoclonal antibodies raised against the classical strain (Samy et al. 2020). Moreover, typical European vvIBDV strains, classical strains, Australian-like strains, and reassortant viruses are still frequently detected in Egypt (Metwally et al. 2009; Samy et al. 2020; Mawgod, Arafa, and Hussein 2014). Over the past few months, field observations have documented numerous instances of bursal atrophy, even in the absence of clinical signs or other gross lesions typically associated with vvIBDV, in both vaccinated and non-vaccinated farms. These cases have coincided with an escalation in the severity of concurrent viral and bacterial infections. To investigate this unusual subclinical form of IBDV, we analyzed samples submitted to our laboratory in 2023, as well as those from previous outbreak episodes in 2021 and 2022, primarily using VP2-HVR for genotyping the circulating viruses..

## Material and methods

### 2.1. Sample collection

Bursal fabricius samples were collected from 21 vaccinated broiler farms suspected of IBDV infection based on clinical diagnosis. These farms were located in major poultry-producing governorates in Egypt during the summer of 2023, including Giza, Beheira, Sharkia, Monufia, Minya, and Matrouh **(Figure 1 and Table 1)**. Each farm was considered as a single epidemiological unit. Additionally, samples from 18 Egyptian farms that had experienced IBDV infections, based on clinical diagnosis, and had submitted samples to the national reference laboratory for veterinary quality control in poultry production during IBDV outbreaks in 2021 and 2022, were included in this study.

**Table 1:** location and epidemiological data of Samples analyzed in this study.

|  | SAMPLES NAMES | PROVINCE | AGE (DAYS) | VACCINE (NO OF DOSES) | GENOTYPE | Accession Numbers |
| --- | --- | --- | --- | --- | --- | --- |
| 1 | IBDV-EGY-F427-2023 | Giza | 20 | Yes (1) | nvarIBDV(A2d) | OR465340 |
| 2 | IBDV-EGY-F647-2023 | Giza | 18 | Yes (2) | nvarIBDV(A2d) | OR465342 |
| 3 | IBDV-EGY-F649-1-2023 | Giza | 22 | Yes (1) | nvarIBDV(A2d) | OR465341 |
| 4 | IBDV-EGY-F649-4-2023 | Giza | 22 | Yes (1) | nvarIBDV(A2d) | OR506926 |
| 5 | IBDV-EGY-F649-5-2023 | Giza | 22 | Yes (1) | nvarIBDV(A2d) | OR506928 |
| 6 | IBDV-EGY-F649-6-2023 | Giza | 22 | Yes (1) | Classic Virulent (A1) | OR465302 |
| 7 | IBDV-EGY-F649-9-2023 | Giza | 22 | Yes (1) | nvarIBDV(A2d) | OR465328 |
| 8 | IBDV-EGY-F533-2023 | Matrouh | 21 | Yes (2) | nvarIBDV(A2d) | OR465333 |
| 9 | IBDV-EGY-F535-2023 | Minya | 23 | Yes (2) | nvarIBDV(A2d) | OR465334 |
| 10 | IBDV-EGY-F555-2023 | Beheira | 16 | Yes (2) | nvarIBDV(A2d) | OR465338 |
| 11 | IBDV-EGY-F558-2023 | Beheira | 21 | Yes (1) | nvarIBDV(A2d) | OR465337 |
| 12 | IBDV-EGY-F621-2023 | Sharkia | 24 | Yes (1) | nvarIBDV(A2d) | OR465332 |
| 13 | IBDV-EGY-F658-1-2023 | Giza | 23 | Yes (1) | nvarIBDV(A2d) | OR465324 |
| 14 | IBDV-EGY-F658-3-2023 | Giza | 20 | Yes (1) | nvarIBDV(A2d) | OR465330 |
| **15** | IBDV-EGY-F65 8-5- 2023 | Giza | 18 | Yes (1) | nvarIBDV(A2d) | OR465329 |
| **16** | IBDV-EGY-F671-2023 | Giza | 24 | Yes (1) | nvarIBDV(A2d) | OR465331 |
| **17** | IBDV-EGY-CV65-2023 | Sharkia | 16 | Yes (1) | nvarIBDV(A2d) | OR465335 |
| **18** | IBDV-EGY-CV67-2023 | Monufia | 16 | Yes (1) | nvarIBDV(A2d) | OR465336 |
| **19** | IBDV-EGY-CV75-2023 | Sharkia | 19 | Yes (1) | nvarIBDV(A2d) | OR465343 |
| **20** | IBDV-EGY-CV98-2023 | Beheira | 27 | Yes (1) | nvarIBDV(A2d) | OR465344 |
| **21** | IBDV-EGY-CV99-2023 | Behiera | 22 | Yes (1) | vvIBDV (A3) | OR465321 |

**Figure 1:**
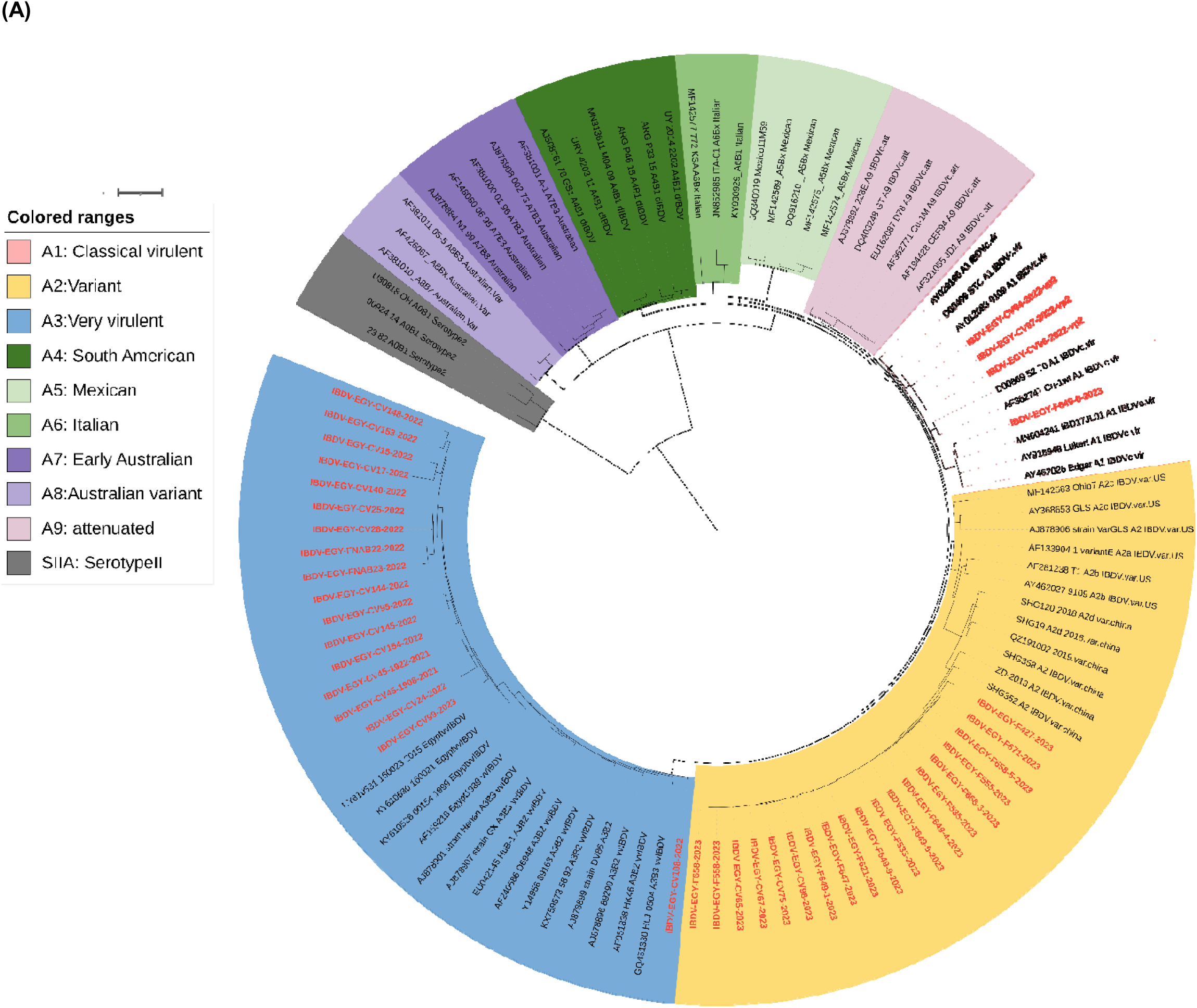
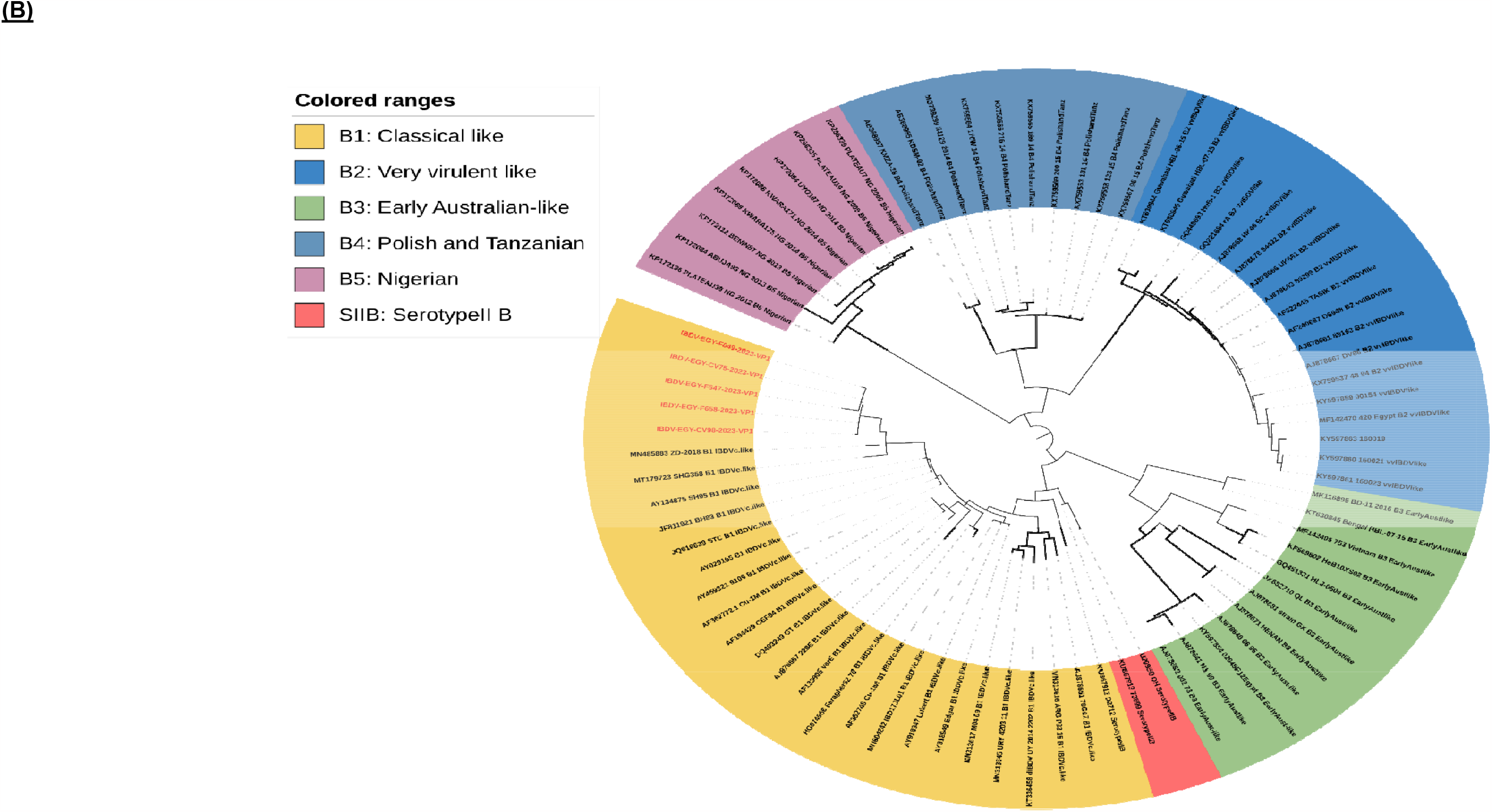
A concise circular phylogenetic analysis based on the VP2-HVR of IBDV segment A (A) and B marker region (B). The tree was generated using a maximum likelihood (ML) phylogenetic tree with 1000 bootstrap replications, using the IQ-TREE program (Nguyen et al. 2015) and the (GTR + F + G4) substitution model, which was determined to be the best-fit model by ModelFinder embedded within IQ-Tree. The tree was annotated in iTOL (https://itol.embl.de) and drawn to scale, with genogroup information displayed in colored circular using scheme proposed by (Gao et al. 2023). Samples tested in the present study colored in red.

### 2.2. Sample preparation

Bursal samples were prepared as previously described (Fan et al. 2019) with some modification. In brief, three to five bursae of Fabricius collected from each farm were weighed, cut using a sterile scalpel and homogenized with an equal mass of phosphate-buffered saline (PBS) supplemented with penicillin and streptomycin. The homogenate was subsequently subjected to three freeze-thaw cycles, followed by centrifugation at 1250xg for 30 minutes. The resulting supernatant was collected and stored at -80°C for further processing.

### 2.3. Viral RNA extraction and qPCR reactions

The prepared Bursal samples were subjected to RNA extraction using Easy Pure® DNA/RNA Extraction Kit (TransGen Biotech, China) according to the manufacturer’s instructions. Reverse transcription and amplification were performed using One Step PrimeScript™ RT-PCR Kit (Takara, Japan) and reaction performed and visualized using Stratagene Mx3000P qPCR System real-time PCR system (Agilent, USA). Primer sets, TaqMan probes and thermal profile used to amplify IBDV VP1 gene were previously described (Moody, Sellers, and Bumstead 2000).

### 2.4. RT-PCR and partial sequencing of VP1 and VP2

The first-strand cDNA synthesis with PCR amplification was performed in a single step using the Easyscript® one-step RT-PCR kit (TransGen Biotech, China) following the manufacturer’s instructions. For the PCR, we used forward primer F673-GTAACAATCACACTGTTCTCAGC and reverse primer R1345 TTATGTCTTAGAAGCCAAATGC for VP2, amplifying a (672bp) segment covering the VP2’s HVR. Additionally, forward primer F320 GAGAATGAGGAGTATGAGACCGA and reverse primer R1150 GAGATCATGAGGTGTGTTGG were used to amplify a (750bp) segment of the VP1 gene. The thermal profile was as follows: 30 min at 50 C, 94 C for 15 min, 40 three-step cycles of 94 C for 30 s, 55 C for 40 s and 72 C for 45 sec; then 72 C for 10 min. After amplification, the PCR products were analyzed by electrophoresis on a 1.5% agarose gel.

The PCR products of the expected sizes were purified from agarose following the manufacturer’s instructions using a QIAquick Gel Extraction Kit (Qiagen GmbH, Hilden, Germany). Sequence reactions were carried out using the Big Dye Terminator Version 3.1 Cycle Sequencing Kit (Applied Biosystems, Foster City, CA, USA). Subsequently, the sequence reactions were purified DyeEx Kits (Qiagen GmbH, Hilden, Germany) and were analyzed using the 3500 XL DNA Analyzer (Applied Biosystems, USA). The retrieved sequences were verified using an online Basic Local Alignment Search Tool (BLAST) (http://www.ncbi.nlm.nih.gov/blast/) to recognize the identity of the sequenced genes.

### 2.5. Phylogenetic and evolutionary analysis

The phylogenetic analysis of different IBDV genotypes used in the present study was based on scheme proposed by (Gao et al. 2023) and confirmed by scehme proposed by (Michel and Jackwood 2017) (Islam et al. 2021) and (Wang et al. 2021) and using samples represent different strains represent variant IBDV subgroups as previously described in (Wang et al. 2022). Multiple sequence alignments were performed using MAFFT online service (Katoh, Rozewicki, and Yamada 2017) and the alignments were visually inspected and manually trimmed using MEGA11 (Tamura, Stecher, and Kumar 2021). The a maximum likelihood (ML) phylogenetic tree with 1000 bootstrap replications was constructed using IQ-TREE program (Nguyen et al. 2015) using (GTR_LJ_+_LJ_F+G4) as a substitution model as the best fit model proposed by ModelFinder embedded within IQ-Tree.

### 2.6. Accession numbers

The partial VP2 sequences for 2021 and 2022 samples were submitted to GenBank under the following accession numbers: OR465303, OR465304, OR465305, OR465307, OR465308, OR465309, OR465310, OR465311, OR465312, OR465313, OR465314, OR465315, OR465316, OR465317, OR465318, OR465319, OR465320 and OR465322. And the partial VP2 sequences for 2023 were submitted to GenBank under the following accession numbers: OR465340, OR465342, OR465341, OR506926, OR506928, OR465302, OR465328, OR465333, OR465334, OR465338, OR465337, OR465332, OR465324, OR465330, OR465329, OR465331, OR465335, OR465336, OR465343, OR465344 and OR465321. While the partial VP1 sequences for 2023 were submitted to GenBank under the following accession numbers: OR465348, OR465350, OR465347, OR465349 and OR465351.

## 3. Results

### 3.1. Field observation and prevalence

Samples were collected from 21 farms for laboratory confirmation of clinical diagnoses over 2023. These farms were located across six governorates, which are major centers for commercial poultry production, mainly in the northern region of Egypt. The sampled farms were broiler production, with birds aged between 2 to 4 weeks, all of which had received vaccinations against infectious bursal disease virus (IBDV), primarily using classical strain-based vaccines (Table 1). According to the farm owners and field veterinarians, historical data and clinical observations revealed that most of farms exhibited bursal atrophy as the predominant lesion. Notably, no other typical IBDV-associated gross lesions, clinical signs, or mortalities were reported. Instead, these farms experienced a significant immunosuppressive effect, leading to reduced vaccine effectiveness against infectious bronchitis and Newcastle disease viruses. Additionally, the severity of infections caused by E. coli and mycoplasma increased, with commonly employed control measures demonstrating limited efficacy. Except for one sample from Behiera governorate, specifically IBDV-EGY-CV98-2023, which exhibited typical IBDV gross lesions that include hemorrhage in bursa, thymus, thigh and breast muscle and was associated with reported mortality **(Table1)**. Regarding the samples delivered to our laboratory during the 2021 and 2022 clinical IBDV episodes, reports indicated typical IBDV gross lesions, symptoms, and reported mortality for the whole outbreaks without data specific for each farms.

### 3.2. Nucleotide sequence and phylogenetic analysis

After confirming the IBDV presence in all samples through real-time PCR, we proceeded with VP2-HVR gene sequencing. Some selected samples also underwent partial VP1 gene sequencing. Phylogenetic analysis was performed using the scheme proposed by (Gao et al. 2023) and confirmed by the schemes proposed by (Islam et al. 2021; Michel and Jackwood 2017; Wang et al. 2021). The results of the phylogenetic analysis based on VP2-HVR revealed that out of the 20 samples collected in 2023, 18 clustered within the A2 genogroup, representing IBDV variants. Within the A2 genotype, four subgroups were identified: A2a, A2b, A2c, and A2d (Gao et al. 2023). All Egyptian samples clustered with the Chinese variant strain in subgroup A2d, but in a separate subgroup (**Figure1, bootstrap value 99%)**. All Egyptian variant strains possess amino acid identities ranging from 97.9% to 98.6% with the recent Chinese strain (HB202201), collected in 2022, and from 95.3% to 96.3% with the older Chinese strain (SHG19), collected in 2018. In terms of identity with U.S. variants, samples in the present study showed 93.2% to 94.6% identity with the variant E strain and 91.2% to 92.8% with the GLS strain. Notably, all Egyptian variant strains in the present study had an identity ranging from 98.1% to 100% between each other.

Additionally,one sample from 2023, along with fifteen samples from 2021 and 2022, clustered within genogroup A3, which represents vvIBDV. Among these, fifteen samples clustered with the Egyptian antigenically atypical vvIBDV isolate 00154 (99323), which were first identified in 1999 (Eterradossi et al. 2004) and have continued to circulate in Egypt over the last two decades (Samy et al. 2020) **(Figure1, bootstrap value 96%)**. These samples displayed an identity ranging from 94% to 98.8% with previous Egyptian isolates from 2015 and 1999. However, only one of the Egyptian vvIBDV strains, namely IBDV-EGY-CV108-2022, did not cluster with the Egyptian vvIBDV group and was closer to the typical European vvIBDV strain (89163) with 98.6% identity. Furthermore, one sample from 2023 and three samples from 2022 clustered with classical virulent IBDV strains within genogroup A1. These samples showed high identity with strains used in vaccines CEVAC IBDL and Bursine, ranging from 98.1% to 100%.

The deduced amino acid sequences of VP2 for the studied samples were aligned with reference samples representing each genogroup. For genogroup A2, samples aligned with Chinese variants belonging to subgroup A2d, and US variants belonging to subgroups A2a, A2b, and A2c. results reveald with except for one sample, all Egyptian variant samples shared the same characteristic amino acids as the reference variant strain Variant E, including 213N, 222T, 242V, 249K, 253Q, 279N, 284A, 286I, 294L, 318D, 323E, and 330S. However, one sample, namely IBDV-EGY-F649-4-2023, possessed 318G, resembling the GLS strain at this residue. Conversely, all Egyptian samples exhibited three conserved amino acid residues, namely 221K, 252I, and 299S, which are unique to Chinese variant IBDVs. Interestingly, all Egyptian variant samples displayed a mutation at residue A321V, which is distinct for Egyptian strains compared to both Chinese and US variants. This mutation is located in major hydrophilic peak B and represents the most exposed part of capsid VP2 **(Figure 2)**.

**Figure 2:**
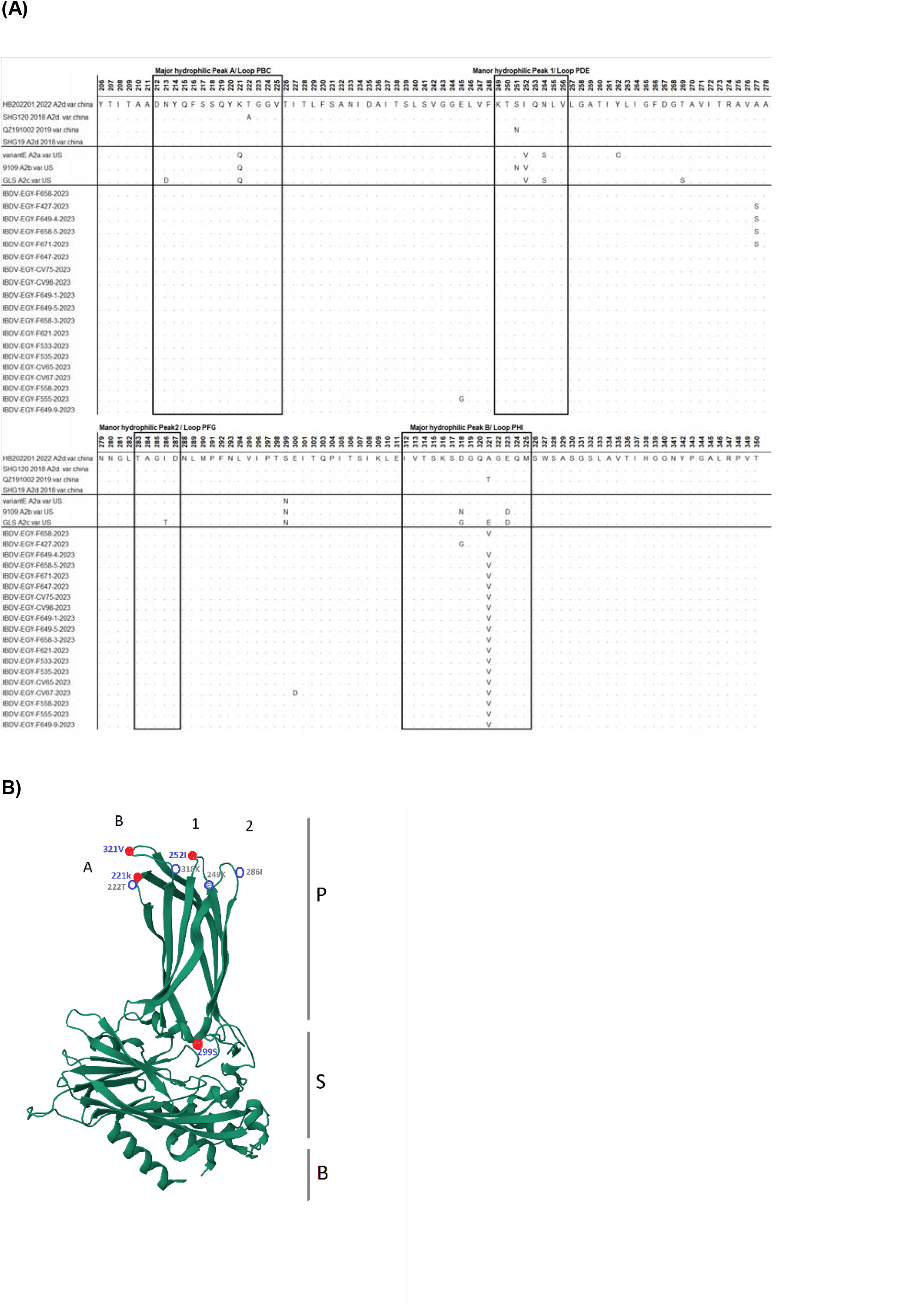
Amino acid alignment of VP2 HVR for the studied Egyptian new variant samples in comparison to US and Chinese variants. Amino acids surrounded by boxes indicate the most exposed peaks of the VP2 P domain (A). The three-dimensional model of IBDV VP2 (PDB-7VRP) was retrieved from the Protein Data Bank to show the localization of amino acids in different VP2 domains (base (B), shell (S), and c projection (P) domains). Close red circles refer to amino acid mutations specific to the new variant with 321V, whi h is only recorded in the Egyptian samples, while blue open circles refer to standard mutations in IBDV variant strains (B).

While for Genogroup A3, samples aligned with typical Europian vvIBDV (89163) and Egyptian antigenically atypical vvIBDV isolate 00154 (99323), which were isolated 1999 and two Egytian vvIBDV strains that represent two subgroups of the Egyptian vvIBDV strains that cocirculating in the field during 2015 survey (Samy et al. 2020), the results revealed presence of 222A, 256I, 294I and 299S residues that considered as markers of vvIBDV in VP2. Interstingly sample IBDV-EGY-CV108-2022, showed identical amino acid sequence of VP2-HVR with typical European vvIBDV strain (89163) with except D279D that reported in Europian genetic reassortant IBDVs with very virulent VP2 (Mató et al. 2020) and I272T. In relation to the atypical vvIBDV strain first identified in global surveillance in 1999, known as 99323, it possesses three unusual mutations: Y220F located in major hydrophilic peak A, G254S in minor hydrophobic peak 1, and A321T in major hydrophilic peak B. These three mutations were responsible for a massive change in the virus’s antigenicity. During the outbreak episode in 2015, two genetic and antigenic patterns were recognized. The first pattern included the three mutations, and this pattern was referred to as strain 160021. The second pattern had two out of the three characteristic mutations, and the residue at 321 resembled that of the European vvIBDV, referred to as strain 160023 **(Figure 3)**. It is noteworthy that strains with the three mutations reacted with 2 out of 8 monoclonal antibodies, while strains with two mutations reacted with 5 out of 8. In contrast, European vvIBDV (89163) reacted with 6 out of 8 monoclonal antibodies raised against classical strains. All the vvIBDV samples detected in the present study possessed only two mutations, Y220F and G254S, but not A321T, indicating antigenicity closer to the European vvIBDV strain.

**Figure 3:**
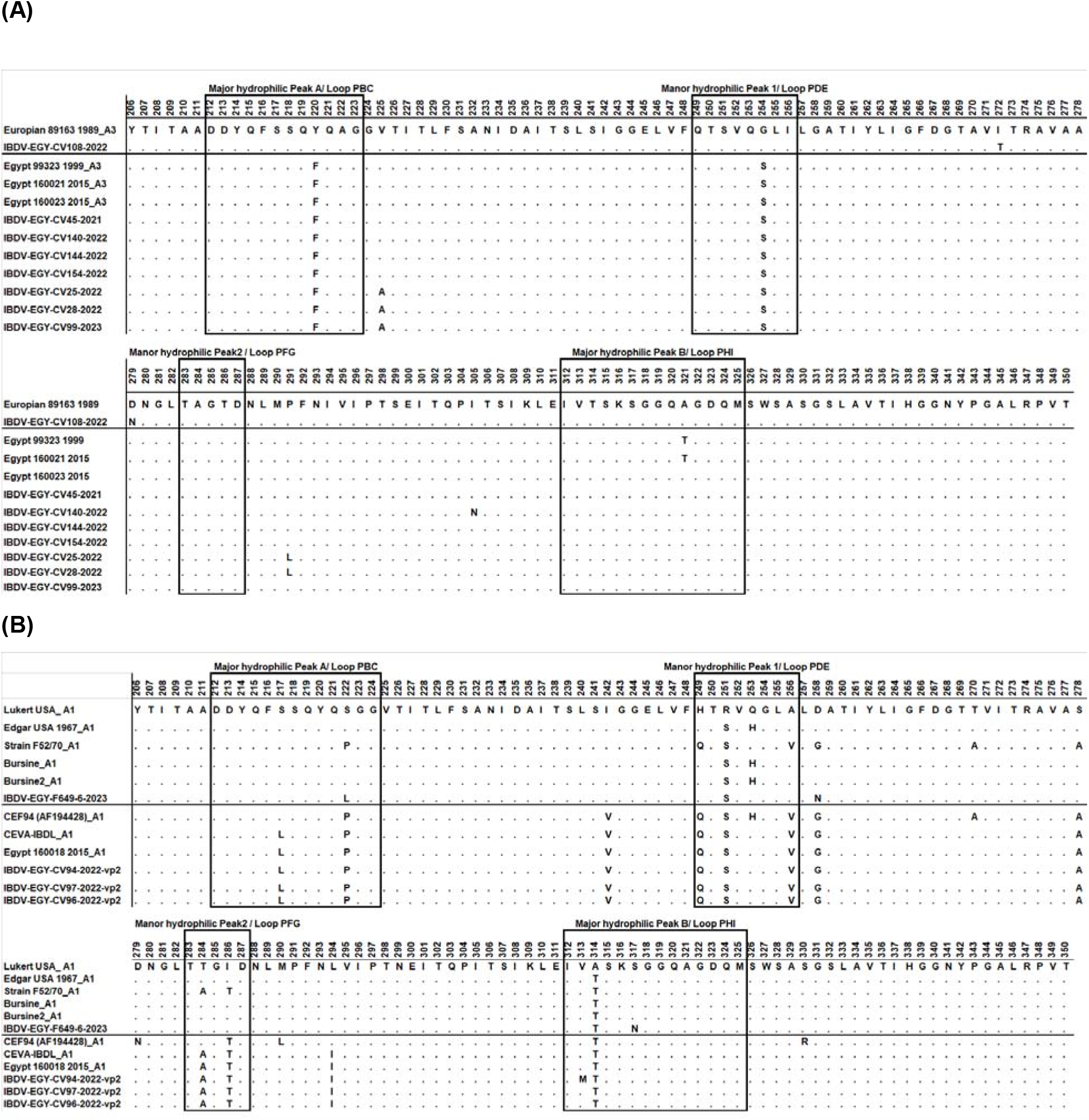
Amino acid alignment of VP2 HVR for the studied Egyptian isolates in comparison with reference sequences. Amino acids surrounded by boxes indicate the most exposed peaks of the VP2 P domain. (A) Represents the amino acid alignment of VP2-HVR for Egyptian vvIBDV in comparison to Typical European vvIBDV (89163) and Egyptian atypical antigenic vvIBDV isolated in 1999 (99323) and 2015 (160021 and 160023). (B) Represents the amino acid alignment of VP2-HVR for Egyptian samples clustered with classical virulent strains in comparison to CEVA IBDL and Bursine strains.

## Discussion

The IBDV was initially recorded in Egypt in the 1970s, coinciding with the introduction of large-scale commercial poultry production (Hassan 2004). By the 1980s, poultry production had significantly increased, driven by substantial government subsidies aimed at promoting egg and poultry consumption. During this period, vvIBDV has been introduced, which was first detected in Europe in 1989 (Zierenberg et al. 2000; El-Batrawy 1990). Despite extensive vaccination efforts, this strain resulted in substantial economic losses (Hassan 2004; Hassan, Afify, and Aly 2002) by end of 1990s the antigenically atypical vvIBDV strain was first isolated that showed to be clinically controlled under experimental condition using life attenuated classic strain based vaccine despite significant changes in the main antigenic domains of VP2 protein (Eterradossi, Gauthier et al. 2004). The antigenically atypical vvIBDV strain has continued to circulate in Egypt over the last two decades, characterized extensive antigenic changes in in the region encoding VP2 projection domain including mutation at positions Y220F and G254S that located in P_BC_ and P_DE_ antigenic domain respectively that detected in all Egyptian vvIBDV, while A321T (P_HI_-Major hydrophilic peak B) frequently detected. vvIBDV has been reported in almost in all major commercial chicken farming regions in Egypt, displaying typical clinical signs and gross lesions including severe haemorrhage in breast and thigh muscles, severely oedematous and haemorrhagic bursa, thymus atrophy, and high mortality rates (Mawgod, Arafa, and Hussein 2014) (Eterradossi et al. 2004). In the present study, we aimed to perform genetic characterization of recently collected samples from farms exhibiting atypical picture of IBDV disease in Egypt where no clinical signs nor post-mortem lesion detected except with atrophied bursa and compare these with samples collected in 2022 and 2021 from farms that displayed typical clinical features of vvIBDV.

The VP2-HVR sequence and partial VP1 (amino acids 216–470) have been identified as phylogenetically representative of the full genome length of IBDV (Le Nouën et al. 2005). The same regions have recently been utilized in several proposed schemes for genotypic classification and nomenclature systems (Michel and Jackwood 2017; Islam et al. 2021; Wang et al. 2021). All of these schemes have reached similar conclusions, indicating the reliability of this system, as reviewed by (Gao et al. 2023). In the present study, we primarily used VP2-HVR to characterize the virus genotypes. Only selected viruses were used to confirm the VP1 using the proposed scheme, as reviewed by (Gao et al. 2023). Our results revealed that eighteen out of twenty samples collected in 2023 clustered with nVarIBDV (A2d) with 97.9 to 98.6% amino acid identity with recent Chinese strain (HB202201). One sample from 2023 clustered with cIBDV (A1 virulent). In contrast, samples from 2021 and 2022 clustered with cIBDV (A1 virulent) and vvIBDV (A3), but none of them belonged to the nVarIBDV. It is worth mentioning that all vvIBDV samples clustered with the antigenically variant vvIBDV that emerged in Egypt in 1999 and has continued to circulate in Egypt over the last two decades, except for one sample (IBDV-EGY-CV108-2022) that clustered with the typical European vvIBDV. Phylogenetic analysis revealed for the first time the prevalence of nVarIBDV in Egypt. The nVarIBDV was first identified in China in 2015 and has been associated with severe economic losses until now. Although it doesn’t cause mortality or morbidity, it does lead to marked bursal atrophy and subsequent severe immunosuppression, increasing the impact of secondary infections by other pathogens. This virus also results in significant economic losses and poses a great challenge to the currently used classical-based vaccines. (Fan et al. 2019; Gao et al. 2023; Wang et al. 2022; Wang et al. 2021) These findings align with the field observations from samples collected in 2023 in the present study. In these cases, all farms displayed bursal atrophy as the primary lesion, with an absence of other typical IBDV-related gross lesions, clinical signs, or mortalities. Instead, there was a pronounced immunosuppressive effect, resulting in reduced vaccine efficacy against infectious bronchitis and Newcastle disease viruses. Furthermore, there was an escalation in the severity of E. coli and mycoplasma infections, and the commonly used control measures showed limited effectiveness. To the best of our knowledge, nVarIBDV has only been reported outside China in Japan (Myint et al. 2021), Korea (Thai et al. 2021) and Malaysia (Aliyu et al. 2021)). This limited spread may be attributed to the virus’s subclinical nature, allowing it to circulate undetected. It’s possible that the virus was directly introduced to Egypt from its original source. This explanation could also clarify the presence of European-like vvIBDV, which is uncommon in Egypt and appears to have been replaced by antigenically modified vvIBDV since 1999 (Samy et al. 2020; Mawgod, Arafa, and Hussein 2014).

The alignment of deduced amino acid sequences for VP2 revealed that the eighteen samples belonging to nVarIBDV exhibit typical residues found in variant IBDV, including 222T, 249K, 286I, and 318D (Jackwood 2012). Additionally, these Egyptian strains have the three conserved amino acid residues that are only found in Chinese variant IBDVs, namely 217K, 252I, and 299S (Fan et al. 2019). Among of them mutation at residues 222 and 249 impacting the virus replication and virulence, while immune escaping associated with mutation at residues 222, 249, 286 and 318 as reviewed by (Gao et al. 2023). However, all Egyptian strains showed a mutation at position 321 (321V), with the exception of one virus that had 318G instead. The mutation at position 318 is associated with immune escape, while the mutation at position 321 critically modifies antigenicity and impacts virulence (Letzel et al. 2007; Escaffre et al. 2013).). It’s worth mentioning that the amino acid at position 321 represents the most exposed part of the capsid protein VP2, and changes in this position have been frequently reported in the antigenically atypical vvIBDV that has been circulating in Egypt since 1999 (Eterradossi et al. 2004) Such changes have been sufficient to induce a dramatic alteration in antigenicity (Samy et al. 2020). Epidemiologically, samples with 318G have only been detected in Giza governorate, while samples with 321V are widely prevalent and have been found in Giza, Beheira, Minya, Matrouh, Sharkia, and Monufia. This indicates the predominance of the 321V genotype and underscores the importance of considering this in any control strategy. It’s important to note that upon examining the sequencing chromatograms of Sanger sequencing files, there were no subpeaks in any of the variant samples at these residues. This rules out the hypothesis of the presence of subpopulations with different sequences from the same genotype or from different genotypes, indicating the co-circulation of both phenotypes in the same governorate. Further sequence epidemiology and laboratory validation of the data are needed to establish a correlation between genotypes and monoclonal antibody reactivity to fully understand the prevalence and impact of these changes.

In the alignment of deduced amino acid sequences for VP2-HVR of cIBDV, there’s a typical amino acid similarity with the commonly used vaccine strain in Egypt. However, in the case of vvIBDV, a careful examination of the sequencing chromatograms revealed the presence of subpeaks in nucleotide sequences associated with changes in residues, particularly in major hydrophilic peak A. This is mainly due to residues like 222, which impact antigenicity, virus replication, and virulence, as discussed previously. Because we couldn’t obtain a pure isolate from these sequences, we decided to exclude them from the deduced amino acid sequences analysis. Among the remaining 9 samples, 8 of them exhibit the typical residues found in the Egyptian antigenically atypical vvIBDV, namely Y220F and G254S. These residues are located in major hydrophilic peak A and manor hydrophobic peak 1, respectively. They are associated with a loss of reactivity to 3 out of 8 monoclonal antibodies raised against the classical IBDV strain, compared to a loss of reactivity to 2 out of 8 for European vvIBDV.(Samy et al. 2020). None of the vvIBDV samples in this study have the A321T residue, which was detected in the first Egyptian antigenically atypical vvIBDV (99323) and co-circulated with the current vvIBDV genotype in 2015, even in the same governorate. This mutation was solely associated with a dramatic change in antigenicity (Eterradossi et al. 2004; Samy et al. 2020; Escaffre et al. 2013; Letzel et al. 2007). Interestingly, one sample (CV108) exhibited a typical amino acid sequence in its major antigenic peaks, resembling European vvIBDV strains. However, it had a change of Asp to Asn at position 279, a mutation previously observed in some European reassortant vvIBDV strains (Mató et al. 2020). This change at residue 279 is linked to adaptation to cell culture and is considered a marker of lower pathogenicity, often associated with subclinical variants and vaccine strains (Noor et al. 2014).

## Conclusion

The present study represents the first report of the emergence of nVarIBDV circulating subclinically in Egypt. The Egyptian variant samples cluster with Chinese nVarIBDV (A2d), all of which exhibit typical residues found in variant IBDV, as well as unique amino acid residues specific to Chinese variant IBDVs. Additionally, they display a unique mutation at position 321, which is the most exposed part of the VP2 capsid. However, the detection of vvIBDV genotypes in only one sample in 2023 does not necessarily imply that nVarIBDV has become the predominant strain. This study has a limited scope and is intended to report the incidence of nVarIBDV in Egypt. Field observations suggest that the subclinical form of IBDV is becoming predominant, while cases of the typical vvIBDV have declined in 2023 compared to previous IBDV outbreak episodes in Egypt. To substantiate this claim, more samples are required from different localities and various production systems, including backyard farming. Furthermore, retrospective sequence analysis indicates that vvIBDV circulating in Egypt still retains its antigenically atypical features, with no detection of A321T, which was associated with significant antigenic changes. It’s worth noting that Egypt has heavily relied on vaccination as a cornerstone in combating IBDV. The current situation highlights two critical points: firstly, vaccines may need revision to ensure protection against the circulating nVarIBDV in Egypt, and secondly, the widespread prevalence of the virus across a large geographic area within a short time suggests potential issues with biosecurity measures that allow virus spread between production units or the introduction of massive infections from outside Egypt. Both hypotheses emphasize the need to review and improve existing biosecurity measures.

## Acknowledgement

We would like to thank farm owners and field veterinarians for their cooperation in sampling, reporting history, clinical signs, and lesions. Special thanks for insightful discussions on clinical observations during current and past IBDV outbreaks. The authors express appreciation to Dr. Ahmed M. El Kady (MEVAC) and Eng. Ahmed M. Badawy (VaccineValley) for supporting AHRI in sampling. The NLQP-AHRI, a governmental institute, receives funding for diagnosis and research from the Agriculture Research Center, Ministry of Agriculture, and Land Reclamation, Egypt. Ahmed Samy is currently supported by the Bill and Melinda Gates Foundation [108704-001] and BBS/E/I/00007038.

## References

Aliyu, Hayatuddeen Bako, Mohd Hair-Bejo, Abdul Rahman Omar, and Aini Ideris. 2021. ‘Genetic diversity of recent infectious bursal disease viruses isolated from vaccinated poultry flocks in Malaysia’, Frontiers in veterinary science, 8: 643976.

Burkhardt, E, and H Müller. 1987. ‘Susceptibility of chicken blood lymphoblasts and monocytes to infectious bursal disease virus (IBDV)’, Archives of virology, 94: 297–303.

Cosgrove, AS. 1962. ‘An apparently new disease of chickens: avian nephrosis’, Avian diseases, 6: 385–89.

El-Batrawy, A. 1990. ‘Studies on Severe Outbreaks of Infectious Bursal Disease’, 2nd Scientific Conference of the Egyptian Veterinary Poultry Association, (Egyptian Veterinary Poultry Association, Cairo): 239–52.

Escaffre, Olivier, Cyril L. Nouën, Michel Amelot, Xavier Ambroggio, Kristen M. Ogden, Olivier Guionie, Didier Toquin, Hermann Müller, Mohammed R. Islam, and Nicolas Eterradossi. 2013. ‘Both Genome Segments Contribute to the Pathogenicity of Very Virulent Infectious Bursal Disease Virus’, Journal of Virology, 87: 2767–80.

Eterradossi, N, JP Picault,P Drouin, Michèle Guittet, Rolande L’Hospitalier, and G Bennejean. 1992. ‘Pathogenicity and preliminary antigenic characterization of six infectious bursal disease virus strains isolated in France from acute outbreaks’, Journal of Veterinary Medicine, Series B, 39: 683–91.

Eterradossi, N, D Toquin, G Rivallan, and M Guittet. 1997. ‘Modified activity of a VP2-located neutralizing epitope on various vaccine, pathogenic and hypervirulent strains of infectious bursal disease virus’, Archives of virology, 142: 255–70.

Eterradossi, N., C. Gauthier, I. Reda, S. Comte, G. Rivallan, D. Toquin, C. D. Boisséson, J. Lamandé, V. Jestin, Y. Morin, C. Cazaban, and P. M. Borne. 2004. ‘Extensive antigenic changes in an atypical isolate of very virulent infectious bursal disease virus and experimental clinical control of this virus with an antigenically classical live vaccine’, Avian Pathology, 33: 423–31.

Eterradossi, Nicolas, and Yehia M. Saif. 2020. ‘Infectious Bursal Disease.’ in, Diseases of Poultry.

Fan, Linjin, Tiantian Wu, Altaf Hussain, Yulong Gao, Xianying Zeng, Yulong Wang, Li Gao, Kai Li, Yongqiang Wang, Changjun Liu, Hongyu Cui, Qing Pan, Yanping Zhang, Yufeng Liu, Hongjiang He, Xiaomei Wang, and Xiaole Qi. 2019. ‘Novel variant strains of infectious bursal disease virus isolated in China’, Veterinary Microbiology, 230: 212–20.

Gao, Hui, Yongqiang Wang, Li Gao, and Shijun J Zheng. 2023. ‘Genetic Insight into the Interaction of IBDV with Host—A Clue to the Development of Novel IBDV Vaccines’, International Journal of Molecular Sciences, 24: 8255.

Hassan, M. K. 2004. ‘Very Virulent Infectious Bursal Disease Virus in Egypt: Epidemiology, Isolation and Immunogenicity of Classic Vaccine’, Veterinary Research Communications, 28: 347–56.

Hassan, M. K., M. Afify, and M. M. Aly. 2002. ‘Susceptibility of vaccinated and unvaccinated Egyptian chickens to very virulent infectious bursal disease virus’, Avian Pathology, 31: 149–56.

Huang, Xuewei, Wei Liu, Junyan Zhang, Zengsu Liu, Meng Wang, Li Wang, Han Zhou, Yanping Jiang, Wen Cui, Xinyuan Qiao, Yigang Xu, Yijing Li, and Lijie Tang. 2021. ‘Very virulent infectious bursal disease virus-induced immune injury is involved in inflammation, apoptosis, and inflammatory cytokines imbalance in the bursa of fabricius’, Developmental & Comparative Immunology, 114: 103839.

Islam, M. R., M. Nooruzzaman, T. Rahman, T. T. Mumu, M. M. Rahman, E. H. Chowdhury, N. Eterradossi, and H. Müller. 2021. ‘A unified genotypic classification of infectious bursal disease virus based on both genome segments’, Avian Pathol, 50: 190–206.

Jackwood, D. J., B. Sreedevi, L. J. LeFever, and S. E. Sommer-Wagner. 2008. ‘Studies on naturally occurring infectious bursal disease viruses suggest that a single amino acid substitution at position 253 in VP2 increases pathogenicity’, Virology, 377: 110–16.

Jackwood, Daral J. 2012. ‘Molecular epidemiologic evidence of homologous recombination in infectious bursal disease viruses’, Avian diseases, 56: 574–77.

Katoh, Kazutaka, John Rozewicki, and Kazunori D Yamada. 2017. ‘MAFFT online service: multiple sequence alignment, interactive sequence choice and visualization’, Briefings in Bioinformatics, 20: 1160–66.

Le Nouën, C., G. Rivallan, D. Toquin, and N. Eterradossi. 2005. ‘Significance of the genetic relationships deduced from partial nucleotide sequencing of infectious bursal disease virus genome segments A or B’, Archives of virology, 150: 313–25.

Letzel, Tobias, Fasseli Coulibaly, Felix A. Rey, Bernard Delmas, Erik Jagt, Adriaan A. M. W. van Loon, and Egbert Mundt. 2007. ‘Molecular and Structural Bases for the Antigenicity of VP2 of Infectious Bursal Disease Virus’, Journal of Virology, 81: 12827–35.

Mató, Tamás, Tímea Tatár-Kis, Balázs Felföldi, Desirée S. Jansson, Zalán Homonnay, Krisztián Bányai, and Vilmos Palya. 2020. ‘Occurrence and spread of a reassortant very virulent genotype of infectious bursal disease virus with altered VP2 amino acid profile and pathogenicity in some European countries’, Veterinary Microbiology, 245: 108663.

Mawgod, Sara Abdel, Abdel Satar Arafa, and Hussein A. Hussein. 2014. ‘Molecular genotyping of the infectious bursal disease virus (IBDV) isolated from Broiler Flocks in Egypt’, International Journal of Veterinary Science and Medicine, 2: 46–52.

Metwally, A. M., A. A. Yousif, I. B. Shaheed, W. A. Mohammed, A. M. Samy, and I. M. Reda. 2009. ‘Reemergence of very virulent IBDV in Egypt’, International Journal of Virology, 5: 1–17.

Michel, L. O., and D. J. Jackwood. 2017. ‘Classification of infectious bursal disease virus into genogroups’, Arch Virol, 162: 3661–70.

Moody, Adrian, Scott Sellers, and Nat Bumstead. 2000. ‘Measuring infectious bursal disease virus RNA in blood by multiplex real-time quantitative RT-PCR’, Journal of Virological Methods, 85: 55–64.

Müller, H. 1986. ‘Replication of infectious bursal disease virus in lymphoid cells’, Archives of virology, 87: 191–203.

Müller, H, and H Becht. 1982. ‘Biosynthesis of virus-specific proteins in cells infected with infectious bursal disease virus and their significance as structural elements for infectious virus and incomplete particles’, Journal of Virology, 44: 384–92.

Müller, Hermann, Md Rafiqul Islam, and Rüdiger Raue. 2003. ‘Research on infectious bursal disease—the past, the present and the future’, Veterinary Microbiology, 97: 153–65.

Myint, Ohnmar, Mathurot Suwanruengsri, Kenji Araki, Uda Zahli Izzati, Apisit Pornthummawat, Phawut Nueangphuet, Naoyuki Fuke, Takuya Hirai, Daral J Jackwood, and Ryoji Yamaguchi. 2021. ‘Bursa atrophy at 28 days old caused by variant infectious bursal disease virus has a negative economic impact on broiler farms in Japan’, Avian Pathology, 50: 6–17.

Nguyen, Lam-Tung, Heiko A Schmidt, Arndt Von Haeseler, and Bui Quang Minh. 2015. ‘IQ-TREE: a fast and effective stochastic algorithm for estimating maximum-likelihood phylogenies’, Molecular Biology and Evolution, 32: 268–74.

Noor, M, MS Mahmud, PR Ghose, U Roy, M Nooruzzaman, EH Chowdhury, PM Das, MR Islam, and H Müller. 2014. ‘Further evidence for the association of distinct amino acid residues with in vitro and in vivo growth of infectious bursal disease virus’, Archives of virology, 159: 701–09.

Raja, P, TMA Senthilkumar, M Parthiban, A Thangavelu, A Mangala Gowri, A Palanisammi, and K Kumanan. 2016. ‘Complete genome sequence analysis of a naturally reassorted infectious bursal disease virus from India’, Genome Announcements, 4: 10.1128/genomea. 00709-16.

Samy, Ahmed, Céline Courtillon, François-Xavier Briand, Mohamed Khalifa, Abdullah Selim, Ahmed Hegazy, Nicolas Eterradossi, and Sébastien M Soubies. 2020. ‘Continuous circulation of an antigenically modified very virulent infectious bursal disease virus for fifteen years in Egypt’, Infection, Genetics and Evolution, 78: 104099.

Tamura, Koichiro, Glen Stecher, and Sudhir Kumar. 2021. ‘MEGA11: Molecular Evolutionary Genetics Analysis Version 11’, Molecular Biology and Evolution, 38: 3022–27.

Thai, Tuyet Ngan, Il Jang, Hyun-A. Kim, Hyun-Soo Kim, Yong-Kuk Kwon, and Hye-Ryoung Kim. 2021. ‘Characterization of antigenic variant infectious bursal disease virus strains identified in South Korea’, Avian Pathology, 50: 174–81.

Vakharia, Vikram N, Junkun He, Basheer Ahamed, and David B Snyder. 1994. ‘Molecular basis of antigenic variation in infectious bursal disease virus’, Virus research, 31: 265–73.

Van den Berg, TP, D Morales, N Eterradossi, G Rivallan, D Toquin, R Raue, K Zierenberg, MF Zhang, YP Zhu, and CQ Wang. 2004. ‘Assessment of genetic, antigenic and pathotypic criteria for the characterization of IBDV strains’, Avian Pathology, 33: 470–76.

Wang, Weiwei, Xiumiao He, Yan Zhang, Yuanzheng Qiao, Jun Shi, Rui Chen, Jinnan Chen, Yanhua Xiang, Zhiyuan Wang, and Guo Chen. 2022. ‘Analysis of the global origin, evolution and transmission dynamics of the emerging novel variant IBDV (A2dB1b): The accumulation of critical aa-residue mutations and commercial trade contributes to the emergence and transmission of novel variants’, Transboundary and Emerging Diseases, 69: e2832–e51.

Wang, Yu-long, Lin-jin Fan, Nan Jiang, Li Gao, Kai Li, Yu-long Gao, Chang-jun Liu, Hong-yu Cui, Qing Pan, Yan-ping Zhang, Xiao-mei Wang, and Xiao-le Qi. 2021. ‘An improved scheme for infectious bursal disease virus genotype classification based on both genome-segments A and B’, Journal of Integrative Agriculture, 20: 1372–81.

Zierenberg, K., H. Nieper, T. P. Van Den Berg, C. D. Ezeokoli, M. Voß, and H. Müller. 2000. ‘The VP2 variable region of African and German isolates of infectious bursal disease virus: Comparison with very virulent, ‘classical’ virulent, and attenuated tissue culture-adapted strains’, Archives of virology, 145: 113–25.

